# Modeling brain dynamics after tumor resection using The Virtual Brain

**DOI:** 10.1101/752931

**Authors:** Hannelore Aerts, Michael Schirner, Thijs Dhollander, Ben Jeurissen, Eric Achten, Dirk Van Roost, Petra Ritter, Daniele Marinazzo

## Abstract

Brain tumor patients scheduled for tumor resection often face significant uncertainty, as the outcome of neurosurgery is difficult to predict at the individual patient level. Recently, computational modeling of brain activity using so-called brain network models has been introduced as a promising tool for this purpose. However, brain network models first have to be validated, before they can be used to predict brain dynamics. In prior work, we optimized individual brain network model parameters to maximize the fit with empirical brain activity. In this study, we extend this line of research by examining the stability of fitted parameters before and after tumor resection, and compare it with baseline parameter variability using data from healthy control subjects. Based on these findings, we perform the first “virtual neurosurgery” analyses to evaluate the potential of brain network modeling in predicting brain dynamics after tumor resection.

We find that brain network model parameters are relatively stable over time in brain tumor patients who underwent tumor resection, compared with baseline variability in healthy control subjects. In addition, we identify several robust associations between individually optimized model parameters, structural network topology and cognitive performance from pre-to post-operative assessment. Concerning the virtual neurosurgery analyses, we obtain promising results in some patients, whereas the predictive accuracy of the currently applied model is poor in others. These findings reveal interesting avenues for future research, as well as important limitations that warrant further investigation.

## 1. Introduction

Many brain tumor patients undergoing neurosurgery face significant uncertainty regarding the outcome of surgery. Although average neurosurgical outcomes for patient cohorts can be predicted with a high degree of accuracy (Emblem et al., 2009, 2015; Senders et al., 2017), the heterogeneity of brain tumors complicates predictions on an individual patient level.

Following methodological advances, several studies have addressed this limitation by applying graph theoretical and machine learning approaches to infer neurosurgical outcome at the individual patient level (for a review see Senders et al., 2018). In particular, several studies have tried to find biomarkers that predict seizure freedom after epilepsy surgery (for example Bonilha et al., 2013, 2015; He et al., 2017; Ji et al., 2015; Morgan et al., 2017; Munsell et al., 2015; Taylor et al., 2018; van Dellen et al., 2014). Others have evaluated machine learning strategies designed to predict survival in glioma (Emblem et al., 2009, 2015) and traumatic brain injury patients (Rughani et al., 2010). Furthermore, one study found that graph measures derived from the pre-surgical functional connectome of patients with temporal lobe epilepsy were able to predict post-surgical cognitive performance scores across different domains (Doucet et al., 2015).

Recently, brain network modeling has also been introduced as a promising tool to simulate neurosurgical outcome (Arsiwalla et al., 2015; Proix, Bartolomei, Guye, & Jirsa, 2017). Brain network modeling techniques implement dynamical models on individual structural brain connectivity networks to simulate subject-specific brain activity (Schirner, McIntosh, Jirsa, Deco, & Ritter, 2018). By virtually lesioning structural connectomes, brain network models may therefore be used as predictive tools to investigate the impact of diverse structural connectivity alterations on brain dynamics, including those purposefully induced by surgery.

For example, a study by Sinha and colleagues (Sinha et al., 2017) investigated surgical outcome in patients undergoing neurosurgery for refractory epilepsy. Specifically, they modeled seizure likelihood per region to identify a highly epileptogenic zone in each patient. According to model predictions, virtual resection of these regions with high seizure likelihood reduced the overall likelihood of seizures, which was confirmed by actual surgical outcomes in the majority of patients (81.3%). Moreover, in patients with poor predicted outcomes, alternative resection sites could be obtained from the model. Furthermore, it has been shown that large-scale brain network models can be used to predict the propagation zone of epileptic activity as determined by stereotactic EEG recordings and clinical expertise (Proix et al., 2017). Importantly, in a follow-up study, they were able to identify the most unstable pathways that support and allow the propagation of seizure activity (Olmi, Petkoski, Guye, Bartolomei, & Jirsa, 2019). Hence, results from this study suggest that selective removal of these unstable connections would be equally effective to render patients seizure free, compared with surgical resection of the entire epileptogenic zone.

The major advantage of brain network modeling is that it produces actual biophysically-oriented models of the brain that go beyond a simple black-box predictor of surgical outcome, potentially making it a useful tool to predict a rich variety of outcomes such as epilepsy status, cognitive performance, functional network integrity and survival. Brain network modeling may thus serve as an important complementary source of information to aid patients and physicians in the process of surgical and medical decision making, by providing estimates of successful and/or adverse outcomes. Furthermore, biologically inspired dynamical models may provide insights into the local dynamics underlying large-scale network topology in health and disease. As such, they may provide an entry point for understanding brain disorders as well as recovery processes after interventions at a causal mechanistic level.

In prior work (Aerts et al., 2018) we investigated brain dynamics before tumor resection in 25 brain tumor patients and 11 healthy control subjects using The Virtual Brain (TVB) (Sanz Leon et al., 2013); an open-source neuroinformatics platform that enables the construction, simulation and analysis of large-scale brain network models. In particular, we optimized model parameters of the Reduced Wong-Wang model (Deco et al., 2014) on an individual basis, after which we compared the fitted parameters between brain tumor patients and healthy control subjects. In addition, we assessed the relations between model parameters, structural network topology and cognitive performance. We found significantly improved prediction accuracy of individual functional connectivity when using individually optimized model parameters, indicating the importance of tuning the model parameters in a subject-specific manner. In addition, local model parameters differed between regions directly affected by a tumor, regions distant from a tumor, and regions in a healthy brain. Lastly, we identified several associations between model parameters, structural network topology and cognitive performance.

In this study, we extend this line of research by examining possible changes in optimized model parameters from pre-to post-operative assessment. To this end, we apply the same procedure as in the pre-operative case to the data acquired several months after each patient’s surgery. To quantify a normal range of baseline variability over time, we also perform parameter optimization on data acquired from healthy control subjects at both time points. After examining the stability of fitted model parameters over time, we use this information to perform the first “virtual neurosurgery” analyses on glioma patients’ pre-operative data, to evaluate the potential of brain network modeling to *predict* brain dynamics after tumor resection.

## 2. Methods

### 2.1 Participants

Patients for this study were recruited with the aim of longitudinal assessment. In particular, data were collected the day before each patient’s tumor resection and again several months after surgery, on the day of patients’ first clinical consultation at the hospital (mean: 7.9 months post-operative; range 5.2–10.7 months post-operative). Patients were included if they were diagnosed with a glioma or meningioma (Fisher, Schwartzbaum, Wrensch, & Wiemels, 2007). Both types of tumors are typically graded according to their malignancy, with grade I tumors being benign, and grade III (for meningioma) or IV (for glioma) being most malignant (Louis et al., 2007). Hereby, malignancy depends on the speed with which the disease evolves, the extent to which the tumor infiltrates healthy brain tissue, and chances of recurrence or progression to higher grades of malignancy.

Patients were recruited from Ghent University Hospital (Belgium) between May 2015 and October 2017. Patients were eligible if they (1) were at least 18 years old, (2) had a supratentorial meningioma (WHO grade I or II) or glioma (WHO grade II or III), (3) were able to complete neuropsychological testing, and (4) were medically approved to undergo MRI investigation. Primary caregivers of the patients were also asked to participate in the study to constitute a group of healthy control subjects that suffer from comparable emotional distress as the patients (Goebel, Von Harscher, & Mehdorn, 2011; Janda et al., 2007). All participants received detailed study information and gave written informed consent prior to study enrollment. This study was approved by the Ethics Committee of Ghent University Hospital.

Out of the 11 glioma patients (mean age 47.5y, *SD* = 11.3y; 36% females), 14 meningioma patients (mean age 60.4y, *SD* = 12.3y; 79% females) and 11 healthy controls (mean age 58.6y, *SD* = 10.3y; 36% females; 10 spouses, 1 adult child) that were included pre-surgically, 7 glioma patients (mean age pre-operatively 50.7y, *SD* = 11.7y; 43% females), 11 meningioma patients (mean age pre-operatively 57.9y, *SD* = 11.0y; 80% females), and 10 control subjects (mean age pre-operatively 59.6, *SD* = 10.3y; 40% females; 9 spouses, 1 adult child) agreed to participate post-operatively. Patient characteristics including follow-up information are described in Table 1.

**Table 1.** Patient characteristics.

| Subject ID | Sex | Age<sup>1</sup> | Handed-ness<sup>1,2</sup> | Tumor lateralization | Tumor location | Tumor histology<sup>3</sup> | Follow-up |
| --- | --- | --- | --- | --- | --- | --- | --- |
| PAT01 | F | 67 | 1.00 | Left | Frontal | Meningioma I | ✓ |
| PAT02 | F | 49 | 1.00 | Right | Frontal | Meningioma I | ✓ |
| PAT03 | F | 60 | 1.00 | Right | Parietal | Meningioma I | ✓ |
| PAT05 | F | 40 | 1.00 | Left | Frontal | Oligo-astrocytoma II | ✓ |
| PAT06 | F | 67 | 1.00 | Bilateral | Frontal | Meningioma I | ✓ |
| PAT07 | M | 55 | -0.16 | Left | Temporal | Ependymoma II | ✓ |
| PAT08 | M | 51 | 1.00 | Left | Frontal | Meningioma I | ✓ |
| PAT10 | F | 77 | 1.00 | Left | Frontal | Meningioma I | ✓ |
| PAT11 | F | 74 | 1.00 | Left | Frontal | Meningioma II | Insufficient data quality |
| PAT13 | F | 40 | -0.68 | Bilateral | Skull base | Meningioma I | ✓ |
| PAT14 | F | 74 | 1.00 | Left | Parietal | Meningioma I | MRI impossible |
| PAT15 | F | 57 | 1.00 | Right | Frontal | Meningioma I | ✓ |
| PAT16 | M | 39 | 1.00 | Right | Fronto-temporal | Anaplastic astrocytoma II-III | ✓ |
| PAT17 | F | 49 | 1.00 | Right | Frontal | Meningioma I | ✓ |
| PAT19 | F | 44 | 1.00 | Right | Frontal skull base | Meningioma I | Drop-out |
| PAT20 | F | 70 | 0.60 | Right | Parietal | Anaplastic astrocytoma III | ✓ |
| PAT22 | M | 53 | 1.00 | Right | Parietal | Oligodendroglioma II | No resection |
| PAT23 | M | 74 | 1.00 | Bilateral | Frontal | Meningioma I | ✓ |
| PAT24 | M | 62 | 1.00 | Left | Frontal | Meningioma I | ✓ |
| PAT25 | F | 46 | 1.00 | Right | Temporal | Glioma II | ✓ |
| PAT26 | M | 61 | 1.00 | Right | Temporal | Anaplastic astrocytoma III | ✓ |
| PAT27 | M | 38 | 1.00 | Left | Frontal | Astrocytoma II | Drop-out |
| PAT28 | M | 44 | 1.00 | Left | Frontal | Oligodendroglioma II | ✓ |
| PAT29 | M | 45 | 0.90 | Left | Frontal | Oligodendroglioma III | End of study |
| PAT31 | F | 31 | 0.90 | Left | Frontal | Oligodendroglioma II | End of study |

<sup>1</sup> Assessed before tumor resection; <sup>2</sup> -1=left-handed, 0=ambidextrous, +1=right-handed; <sup>3</sup> (Oligo-)astrocytoma, ependymoma, and oligodendroglioma are subtypes of glioma tumors.

### 2.2 MRI data acquisition and preprocessing

MRI sequence details and preprocessing procedures for the post-operative data are mostly identical to those that we used before to collect and preprocess the pre-operative data. All details are described in Aerts et al. (2018). In the following sections, we provide a summary of these procedures, as well as an overview of the minor modifications that were applied.

#### 2.2.1 MRI data acquisition

From all participants, three types of MRI scans were obtained using a Siemens 3T Magnetom Trio MRI scanner with a 32-channel head coil. First, a T1-weighted MPRAGE image was acquired (160 slices, TR = 1750 ms, TE = 4.18 ms, field of view = 256 mm, flip angle = 9°, voxel size 1 × 1 × 1 mm, acquisition time of 4:05 min). Next, resting-state functional echo-planar imaging (EPI) data were obtained in an interleaved order (42 slices, TR = 2100ms, TE = 27 ms, field of view = 192 mm, flip angle = 90°, voxel size 3 × 3 × 3 mm, acquisition time of 6:24 min)^1^. During the fMRI scan, participants were instructed to keep their eyes closed and not fall asleep. Finally, multi-shell high-angular resolution diffusion imaging (HARDI) MRI data were acquired (60 slices, TR = 8700 ms, TE = 110 ms, field of view = 240 mm, 102 gradient directions, b-values of 0, 700, 1200, 2800 s/mm^2^, voxel size 2.5 × 2.5 × 2.5 mm, acquisition time of 15:14 min) (Jeurissen, Tournier, Dhollander, Connelly, & Sijbers, 2014). In addition, two diffusion MRI b = 0 s/mm^2^ images were collected with reversed phase-encoding blips to correct susceptibility induced distortions (Andersson, Skare, & Ashburner, 2003).

#### 2.2.2 Preprocessing of T1-weighted anatomical MRI data

High-resolution anatomical images were processed using the default “recon-all” processing pipeline of FreeSurfer (http://surfer.nmr.mgh.harvard.edu), yielding a subject-specific parcellation of each participant’s cortex into 68 regions (Desikan et al., 2006; Fischl et al., 2004). To account for lesion effects in the parcellation, some additional steps were performed depending on the specific tumor type. For meningioma tumors, that were completely resected during neurosurgery, few to no lesion effects were apparent in eight out of ten patients. For these patients, the default processing pipeline was applied. In the other two meningioma patients, residual edema and a resection cavity were observed, respectively. Therefore, we used a procedure similar to the one outlined in Solodkin et al. (2010). Specifically, we first produced an enantiomorphic filling of the lesioned area (Nachev, Coulthard, Jäger, Kennard, & Husain, 2008) using the BCBtoolkit (Foulon et al., 2018), after which the standard FreeSurfer processing pipeline was utilized. For the glioma patients, we used each patient’s pre-operative parcellation scheme, after non-linear registration to their post-operative space (using FSL FNIRT; Andersson, Jenkinson, & Smith, 2007). All registration results were visually verified.

#### 2.2.3 Functional MRI preprocessing

Resting-state fMRI data were preprocessed using FEAT (FMRI Expert Analysis Tool, version 6.00), part of FSL (FMRIB’s Software Library; Jenkinson, Beckmann, Behrens, Woolrich, & Smith, 2012), comprising motion correction, slice-timing correction, non-brain removal, grand-mean intensity normalization and high-pass temporal filtering (100-second high-pass filter). Functional connectivity matrices were then constructed by mapping the FreeSurfer cortical parcellation schemes obtained in the previous step to each subject’s functional MRI data, and calculating the Fisher’s z-transformed Pearson correlation coefficient between all region-wise BOLD time series.

#### 2.2.4 Diffusion MRI preprocessing

For preprocessing and construction of structural connectomes based on the diffusion MRI (dMRI) data, a processing pipeline was used combining FSL (FMRIB’s Software Library; Jenkinson et al., 2012; version 5.0.9) and MRtrix3 (Tournier et al., 2019). Preprocessing steps included correction for various artifacts (noise (Veraart et al., 2016), Gibbs ringing (Kellner, Dhital, Kiselev, & Reisert, 2016), motion and eddy currents (Andersson & Sotiropoulos, 2016), susceptibility induced distortions (Andersson, Skare, & Ashburner, 2003) and bias field inhomogeneities (Zhang, Brady, & Smith, 2001)), registration of subjects’ high-resolution anatomical images to diffusion space (Jenkinson, Bannister, Brady, & Smith, 2002; Jenkinson & Smith, 2001), and segmentation of the anatomical images into gray matter, white matter and cerebrospinal fluid (Zhang et al., 2001). Further, quantitative whole-brain probabilistic tractography was performed using MRtrix3 (Tournier et al., 2019), resulting in 7.5 million streamlines per subject (more details are available in Aerts et al., 2018). Structural connectivity (SC) matrices were then constructed by transforming each individual’s FreeSurfer parcellation scheme to the diffusion MRI data and calculating the number of estimated streamlines between each pair of brain regions. Lastly, we thresholded the SC matrices and normalized structural connections with the same constant scalar as in the pre-operative analyses, to ensure all weights varied between 0 and 1 and were maximally comparable between pre- and post-operative assessment.

### 2.3 Brain network modeling

Procedures for simulating large-scale brain dynamics and optimizing model parameters were also identical to those applied to the pre-operative data, as described in detail in Aerts et al. (2018). Briefly, local dynamics for each of the 68 cortical brain regions were simulated using Reduced Wong-Wang neural mass models (Deco et al., 2014), which faithfully approximate the mean dynamics of interacting populations of excitatory and inhibitory spiking neurons. Subsequently, neural mass models were coupled according to each subject’s tractography-derived structural connectome to generate personalized virtual brain models (Deco et al., 2014; Ritter, Schirner, McIntosh, & Jirsa, 2013; Sanz Leon et al., 2013; Schirner et al., 2018).

To optimize the correspondence between empirical and simulated functional connectivity, subject-specific parameter space explorations were conducted in which the global scaling parameter (G) was varied (0.01 to 3 in steps of 0.015). This parameter rescales each subject’s structural connectivity, which is given by relative values, to yield absolute interaction strengths. For each parameter set, resting-state blood-oxygen-level-dependent (BOLD) time series were generated. Subsequently, functional connectivity matrices were computed by calculating the Fisher’s z-transformed Pearson correlation coefficient between all pairs of simulated BOLD time series. The parameter set that maximized the Pearson correlation between each individual’s simulated and empirical functional connectivity matrix was then selected for further analyses.

In addition, inhibitory synaptic weights (J_i_) – which control the strength of connections from inhibitory to excitatory mass models within each large-scale region *i* – were automatically tuned in each iteration of the parameter space exploration, to clamp the average firing rate at 3 Hz for each excitatory mass model (Deco et al., 2014; Schirner et al., 2018). After simulations, the obtained local inhibitory connection strengths were corrected for their respective region size for further analyses, since the need for local inhibition to balance global excitation depends on the total connection strength a brain region has, which tightly correlates with region size (Aerts et al., 2018). Median J_i_ values (both corrected for region size as well as uncorrected) across the entire brain and across tumor and non-tumor regions in brain tumor patients were then computed per subject. Of note, delineation of tumor and non-tumor regions was based on the pre-operative data. A more detailed description is available in Aerts et al. (2018).

### 2.4 Graph analysis

Post-operative structural network topology was evaluated using the same graph metrics as those applied to the pre-operative structural connectomes (Aerts et al., 2018). Specifically, global efficiency, modularity and participation coefficient were computed using the Brain Connectivity Toolbox (see Rubinov & Sporns (2010) for more details and an in-depth discussion of graph metrics).

### 2.5 Neuropsychological testing

Cognitive performance of all participants was re-assessed after each patient’s tumor resection using the Cambridge Neuropsychological Test Automated Battery (CANTAB®; Cambridge Cognition (2017); All rights reserved; http://www.cambridgecognition.com). The same cognitive tasks were administered as before surgery, again in random order to avoid sequence bias. In particular, the Rapid Visual Information Processing (RVP) task was used to assess sustained attention, the Spatial Span (SSP) task measured working memory capacity, the Reaction Time task (RTI) evaluated mental response speed, and the Stockings of Cambridge (SOC) task assessed planning accuracy.

### 2.6 Accounting for covariates of no immediate interest

Several factors can influence cognitive performance and graph metrics (see for example Bettus et al., 2010; Biswal et al., 2010; Harrison et al., 2008). Therefore, cognitive performance results were corrected for each participant’s level of emotional distress, residual lesion size, age and sex. Likewise, graph metrics were corrected for each subject’s level of emotional distress, residual lesion size, age, sex, handedness, motion during resting-state fMRI acquisition and intensity normalization factor used in dMRI preprocessing. In particular, on the day testing took place, emotional distress was measured using the State-Trait Anxiety Inventory (Spielberger, Gorsuch, Lushene, Vagg, & Jacobs, 1983; Van der Ploeg, 1982). Further, residual lesion volume was calculated as the number of 1 mm^3^ isotropic voxels in the mask that delineated residual lesion tissue, which was drawn manually on the anatomical T1-weighted MRI image. Lastly, handedness was measured using the Edinburgh Handedness Inventory (Oldfield, 1971).

We then constructed linear regression models for every outcome variable (sustained attention, working memory capacity, reaction time, and planning accuracy for cognitive performance; global efficiency, modularity and participation coefficient for graph theory metrics) as a function of these confounders. Residuals of these models were further transformed to z-scores for subsequent analyses using the pre-operative mean and standard deviation of the respective metric in the group of control subjects, for ease of interpretation.

### 2.7 Statistical analyses

First, we compared post-operatively optimized model parameters, cognitive performance scores and graph metrics between glioma patients, meningioma patients and control participants. For these analyses, we used one-way analysis of variance (ANOVA) and Kruskal-Wallis rank sum tests, depending on whether or not the normality assumption held. Next, we computed difference scores between each participant’s pre- and post-surgical model parameters, cognitive performance scores and network topology indices to evaluate whether changes over time were evident using one-sample t-tests. In addition, group differences in the mean and variance of difference scores between pre- and post-operative assessment were examined using one-way ANOVA and Levene’s test for equality of variances. Afterwards, post-surgical optimal model parameters were related to structural network topology and cognitive performance using linear regression. Likewise, difference scores between pre- and post-operatively optimized model parameters were compared with differences in cognitive performance scores and graph metrics over time. Statistical analyses were carried out with R version 3.5.3 (R Core Team, 2018).

### 2.8 Virtual tumor resection

After examining the stability of fitted model parameters over time, this information was used to perform the first virtual neurosurgery analyses, to evaluate the potential of brain network modeling to *predict* brain dynamics after tumor resection. To this end, a procedure similar to the one described by Taylor and colleagues (Taylor et al., 2018) was adopted. In particular, each patient’s actual surgery was mimicked by removing all streamlines from their pre-operative tractogram that intersect the resection mask that was retrospectively derived from the post-operative anatomical MRI data. Since standard tractography algorithms are currently unable to reliably reconstruct white matter streamlines within or in close proximity to tumorous tissue, a dedicated pipeline was developed for this second part of the study. This was of crucial importance to allow simulation of tumor resection procedures, since white matter tracts in the vicinity of the tumor have the highest probability of being removed during neurosurgery. These proof of concept analyses were performed on all glioma patients for which both pre- and post-operative data were available. Virtual neurosurgery was not performed on data from meningioma patients, as these tumors generally do not infiltrate healthy brain tissue and therefore are not represented within the tractogram or structural connectivity matrix. Figure 1 illustrates the procedure used to predict post-surgical brain dynamics after virtual tumor resection, using brain network modeling.

**Figure 1.**
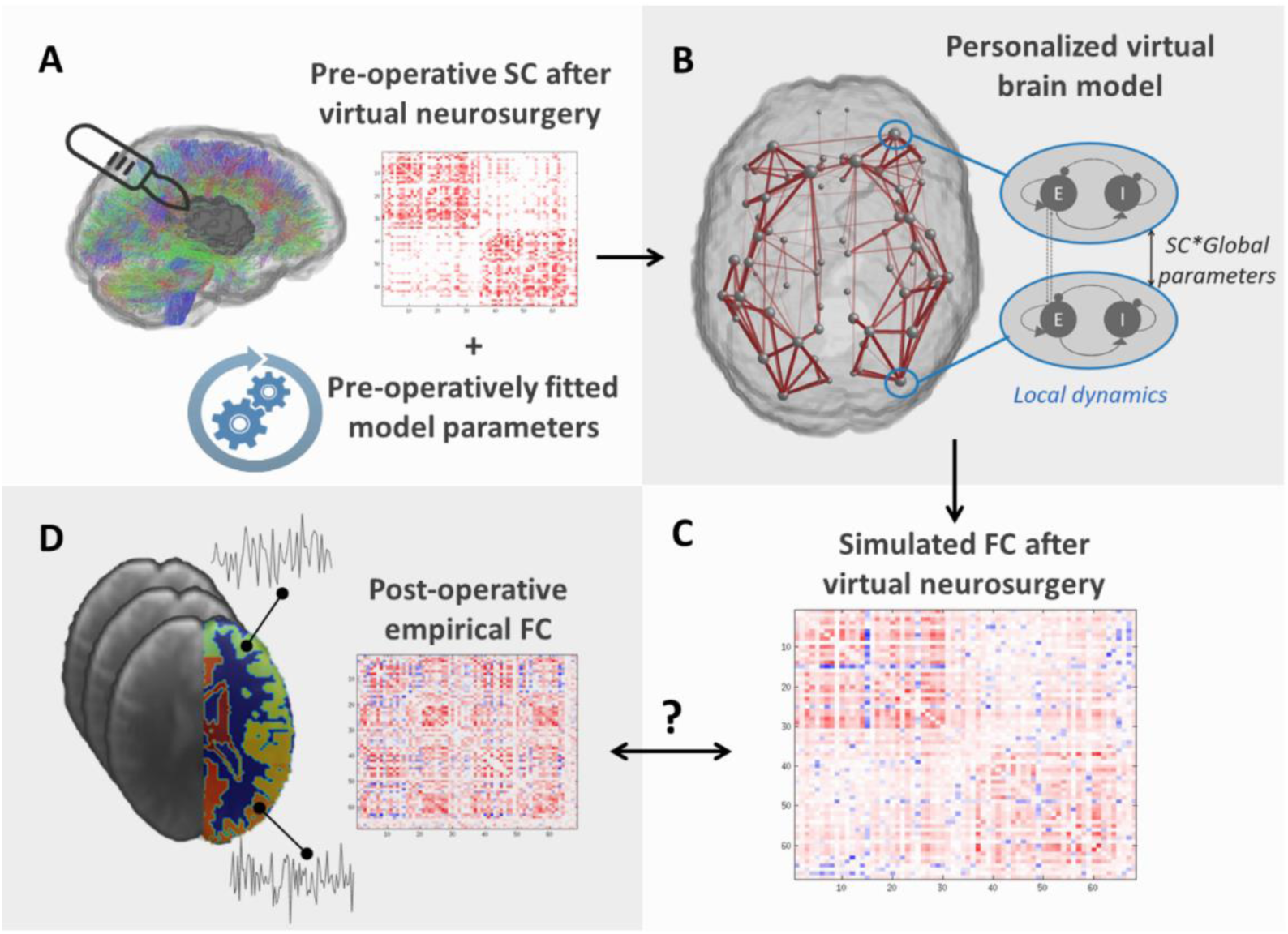
Graphical overview of the procedure to evaluate the potential of brain network modeling after virtual neurosurgery to predict post-surgical brain dynamics. **A:** First, each patient’s pre-operative structural connectome is reconstructed using dMRI whole-brain streamlines tractography, after which actual surgery is mimicked by removing all streamlines that intersect the resection mask that was retrospectively derived from the post-operative anatomical MRI data. Additionally, pre-operatively optimized model parameters, fitted to the subject’s pre-operative functional connectivity, are supplied. **B:** Subsequently, large-scale brain dynamics are simulated on patients’ virtually lesioned structural connectome. **C:** Resulting brain dynamics are transformed to a functional connectivity matrix, representing brain dynamics after virtual neurosurgery. **D:** Simulated brain dynamics are compared to patients’ empirical FC, derived from their post-operative fMRI data that served as ground truth.

#### 2.8.1 Reconstruction of pre-operative tumor structural connectome

Starting from the preprocessed pre-operative dMRI data, a novel technique to model the diffusion signal was applied, named single-shell 3-tissue constrained spherical deconvolution (SS3T-CSD) (Dhollander & Connelly, 2016; Dhollander, Raffelt, & Connelly, 2016), using MRtrix3Tissue (https://3Tissue.github.io), a fork of the MRtrix3 software (Tournier et al., 2019). Investigating the performance of this technique for this purpose, we have recently shown that SS3T-CSD allows to reconstruct white matter streamlines within infiltrative tumors and in immediately adjacent tissue (Aerts, Dhollander, & Marinazzo, 2019).

Based on the resulting white matter fiber orientation distributions (FODs), probabilistic streamlines tractography was performed, using an FOD amplitude threshold of 0.07 (Tournier, Calamante, & Connelly, 2010). Although white matter FODs could be estimated using SS3T-CSD within regions infiltrated by a tumor (Aerts et al., 2019), resulting white matter FOD amplitudes were substantially smaller in tumor regions compared to the rest of the brain. While this likely reflects the smaller portion of space taken up by axons (due to infiltrating tumor tissue) and/or damage to white matter tracts, it does pose a practical challenge to tractography algorithms, which rely on the aforementioned amplitude threshold to determine where and how far tractography may proceed. As detailed in Aerts et al. (2019), we overcame this by gradually reducing the FOD amplitude threshold close to and even more so within the tumor, based on its prior segmentation. Anatomical constraints were also imposed to the generation of streamlines, informed by a segmented tissue image (Smith, Tournier, Calamante, & Connelly, 2012). Since this particular segmentation strategy misclassified tumorous tissue mostly as gray matter, which is then enforced to be an endpoint of white matter streamlines during tractography, a *modified* segmented tissue image was provided. Specifically, the tumor mask was filled with undamaged tissue from homologous regions within the contralateral hemisphere using a non-linear registration approach (Foulon et al., 2018; Nachev et al., 2008), providing an approximation of the patient’s brain anatomy as if the tumor were absent. This “restored” anatomical image was then segmented and used to generate 30 million streamlines connecting pairs of brain regions.

All reconstructed streamlines were filtered to 7.5 million tracts using SIFT (Smith, Tournier, Calamante, & Connelly, 2013) to obtain quantitative streamline counts. Structural connectivity matrices were then constructed by calculating the number of estimated streamlines between any two Desikan-Killiany cortical brain regions, and normalizing all connectivity weights with a (single) constant scalar across subjects to ensure all weights varied between 0 and 1.

#### 2.8.2 Identifying optimal model parameters

Using the newly constructed pre-surgical structural connectome, we redid subject-specific parameter space explorations to identify each patient’s optimal global coupling value that maximized the correspondence between pre-surgical empirical and simulated functional connectivity. To this end, we adopted the procedure as described in section 2.3 (“Brain network modeling”).

#### 2.8.3 Virtual lesioning of the structural connectome

To mimic each patient’s actual neurosurgical procedure retrospectively, tumor resection cavity maps were drawn manually under the supervision of an expert neuroradiologist (E.A.) based on the patient’s post-operative anatomical T1-weighted MRI data. These resection masks were then overlaid onto the patient’s pre-surgical tractogram using non-linear registration, and all connections that intersected the resection cavity mask were removed, similar to the approach outlined in Taylor et al. (2018).

#### 2.8.4 Simulating post-surgical brain dynamics

Using the patient’s pre-surgically optimized model parameters and virtually lesioned structural connectome, we then simulated large-scale brain dynamics with the Reduced Wong Wang model (Deco et al., 2014). Finally, each patient’s predicted functional connectome was compared to their post-operative empirical functional connectome that served as ground truth, by means of link-wise Pearson correlation.

## 2.9 Data and code accessibility

All data and code used for this study is freely available. The data is publicly available at the OpenNeuro website (https://openneuro.org) and on the European Network for Brain Imaging of Tumours (ENBIT) repository (https://www.enbit.ac.uk) under the names “BTC_preop” and “BTC_postop” for the pre- and post-operative data, respectively. The optimized TVB C code can be found at https://github.com/BrainModes/The-Hybrid-Virtual-Brain and all scripts for postprocessing can be found at https://github.com/haerts/The-Virtual-Brain-Tumor-free-Patient.

## 3. Results

### 3.1 Stability of individual model parameters over time

In the first part of this study we examine if and how individually optimized model parameters change after tumor resection. In order to identify normal ranges of variability over time, we perform the same analyses in a group of healthy control subjects. Results are summarized in Table 2 and Figure 2. In particular, the subplots on the left of Figure 2 show the difference scores between pre- and post-operatively optimized model parameters in meningioma patients, glioma patients and control subjects. Differences over time in fitted parameters are shown as deviations from the horizontal line drawn around zero, with positive scores indicating increases in post-operative relative to pre-operative measures, whereas negative scores correspond to decreases after surgery compared with pre-operative levels. The subplots on the right of Figure 2 depict pre-versus post-operatively optimized model parameters at the individual level, shape- and color-coded by group. Here, differences over time in individuals’ fitted parameters are shown as deviations from the main diagonal, with scores above the diagonal indicating increases in post-operative relative to pre-operative measures, whereas measures below the main diagonal correspond to decreases after surgery compared with pre-operative levels.

**Table 2.** Statistical results of analyses on differences between groups and over time of optimized model parameters.

|  | Pre-operative | Post-operative | Difference scores (post-pre) |
| --- | --- | --- | --- |
| **J<sub>tumor</sub>**<br>(corrected<br>for ROI size) | Group differences mean:<br>$\chi^2(2) = 18.95, p < .0001$<br>Group differences variance:<br>$F(2,24) = 153.3, p < .0001$ | Group differences mean:<br>$\chi^2(2) = 15.28, p = 0.0005$<br>Group differences variance:<br>$F(2,24) = 62.07, p < .0001$ | Difference across groups:<br>$t(26) = 1.58, p = 0.13$<br>Group differences mean:<br>$\chi^2(2) = 2.89, p = 0.24$<br>Group differences variance:<br>$F(2,24) = 2.00, p = 0.16$ |
| **J<sub>non-tumor</sub>**<br>(corrected<br>for ROI size) | Group differences mean:<br>$F(2,24) = 4.58, p = 0.0207$ | Group differences mean:<br>$F(2,24) = 7.82, p = 0.0024$ | Difference across groups:<br>$t(26) = 2.84, p = 0.0087$<br>Group differences mean:<br>$F(2,24) = 0.66, p = 0.53$<br>Group differences variance:<br>$F(2,24) = 0.25, p = 0.78$ |
| **J<sub>tumor</sub>**<br>(uncorrected) | Group differences mean:<br>$\chi^2(2) = 7.78, p = 0.0204$<br>Group differences variance:<br>$F(2,24) = 4.12, p = 0.0291$ | Group differences mean:<br>$F(2,24) = 5.76, p = 0.0091$<br>Group differences variance:<br>$F(2,24) = 9.69, p = 0.0008$ | Difference across groups:<br>$t(26) = 0.27, p = 0.79$<br>Group differences mean:<br>$\chi^2(2) = 2.71, p = 0.26$<br>Group differences variance:<br>$F(2,24) = 2.76, p = 0.08$ |
| **J<sub>non-tumor</sub>**<br>(uncorrected) | Group differences mean:<br>$F(2,24) = 1.90, p = 0.17$<br>Group differences variance:<br>$F(2,24) = 0.55, p = 0.58$ | Group differences mean:<br>$F(2,24) = 1.15, p = 0.33$<br>Group differences variance:<br>$F(2,24) = 1.10, p = 0.35$ | Difference across groups:<br>$t(26) = 1.21, p = 0.24$<br>Group differences mean:<br>$F(2,24) = 0.18, p = 0.84$<br>Group differences variance:<br>$F(2,24) = 0.22, p = 0.81$ |
| **G** | Group differences mean:<br>$F(2,24) = 2.23, p = 0.13$ | Group differences mean:<br>$F(2,24) = 0.75, p = 0.48$ | Difference across groups:<br>$t(26) = 1.24, p = 0.23$<br>Group differences mean:<br>$\chi^2(2) = 3.65, p = 0.16$<br>Group differences variance:<br>$F(2,24) = 1.07, p = 0.36$ |

**Figure 2.**
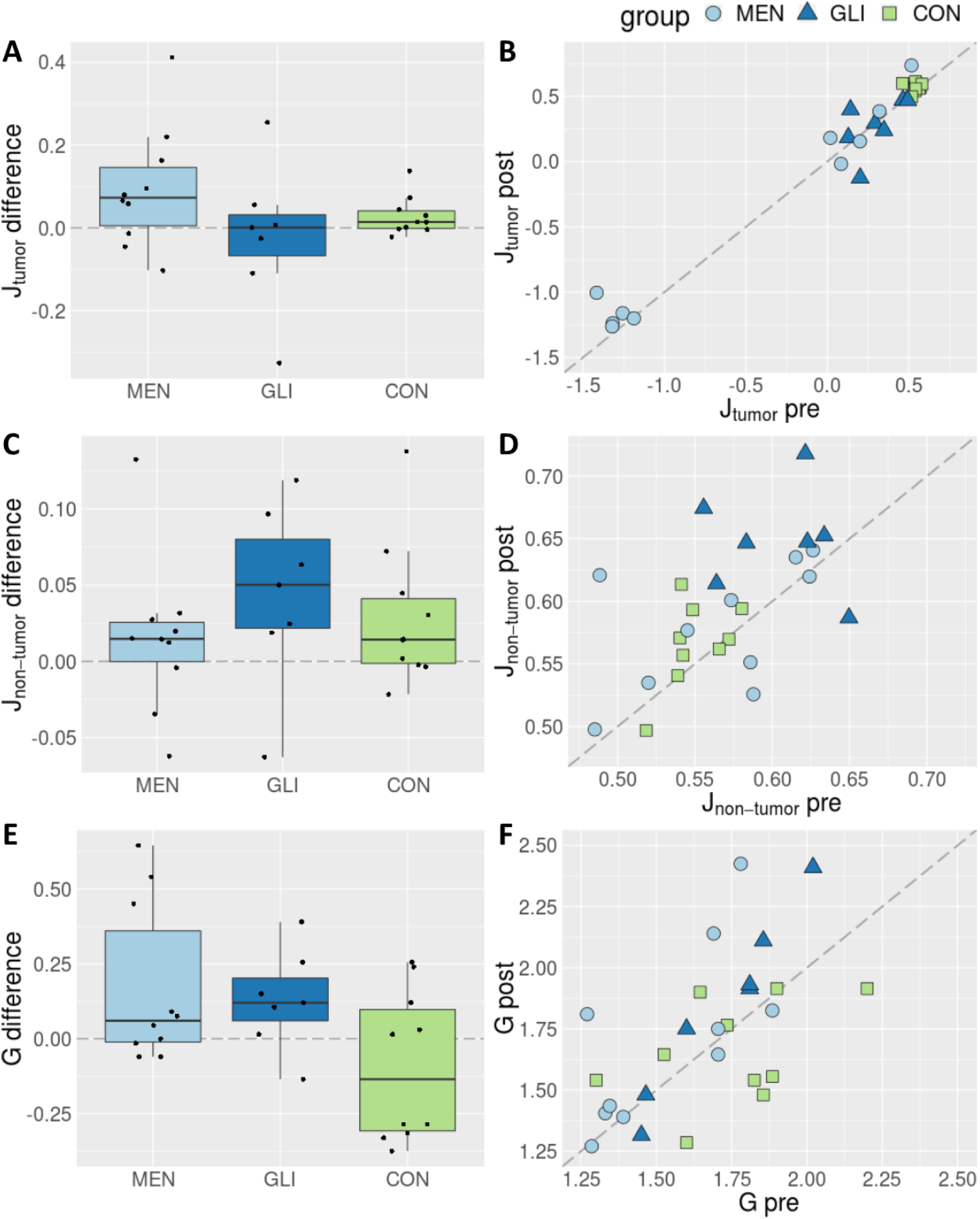
Difference scores between pre- and post-operatively optimized model parameters by group (left) and pre-versus post-operatively optimized individual fitted parameters (right) in meningioma patients (MEN), glioma patients (GLI) and control subjects (CON). **A & B:** Median local inhibitory connection strengths (corrected for region size) across tumor regions in meningioma and glioma patients, and across entire brain in healthy controls; **C & D:** Median local inhibitory connection strengths (corrected for region size) across non-tumor regions in meningioma and glioma patients, and across entire brain in healthy controls, and; **E & F:** global scaling parameter.

Specifically, Figure 2A and 2B depict the median local inhibitory connection strengths across tumor regions in meningioma and glioma patients, and across the entire brain in healthy controls (after correcting for region size). Changes over time in local inhibitory connection strengths are not statistically significant, nor do they differ significantly between groups. Nevertheless, local inhibitory connection strengths are much lower and more variable in tumor regions compared to those in healthy brains, before as well as after tumor resection.

Further, results reveal differences in median local inhibitory connection strengths (corrected for region size) across non-tumor regions in tumor patients relative to those in healthy brains (Figure 2C and 2D). This is the case both before patients’ tumor resection, as well as after surgery. Moreover, we observe a general increase in participants’ median local inhibitory connection strengths after surgery compared to their pre-surgical levels, although changes over time do not differ significantly between groups.

Of note, without correcting for region size, we also find significant differences in median local inhibitory connection strengths between tumor and healthy brain regions. However, without correction for region size, brain tumor patients show higher levels of feedback inhibition compared to controls, whereas the opposite trend was found when correcting for region size (see Supplementary Figure 1A and 1B). Changes over time in local inhibitory connection strengths are also not statistically significant, and do not differ significantly between groups. In contrast, no significant differences are apparent in median local inhibitory connection strengths between non-tumor and healthy brain regions without correction for region size (Supplementary Figure 1C and 1D).

Finally, we find no statistically significant group differences nor effects over time in the global scaling parameter (Figure 2E and 2F).

### 3.2 Stability of structural network topology and cognitive performance over time

Before relating optimized model parameters to cognitive performance scores and structural network topology metrics, we examine whether changes in these predictor variables can be observed over time, or whether post-operative group differences are apparent. Statistical results are summarized in Table 3, and Supplementary Figure 2 and 3 provide a visual overview of pre- and post-operative cognitive performance and graph metrics (z-scores), respectively.

**Table 3.**
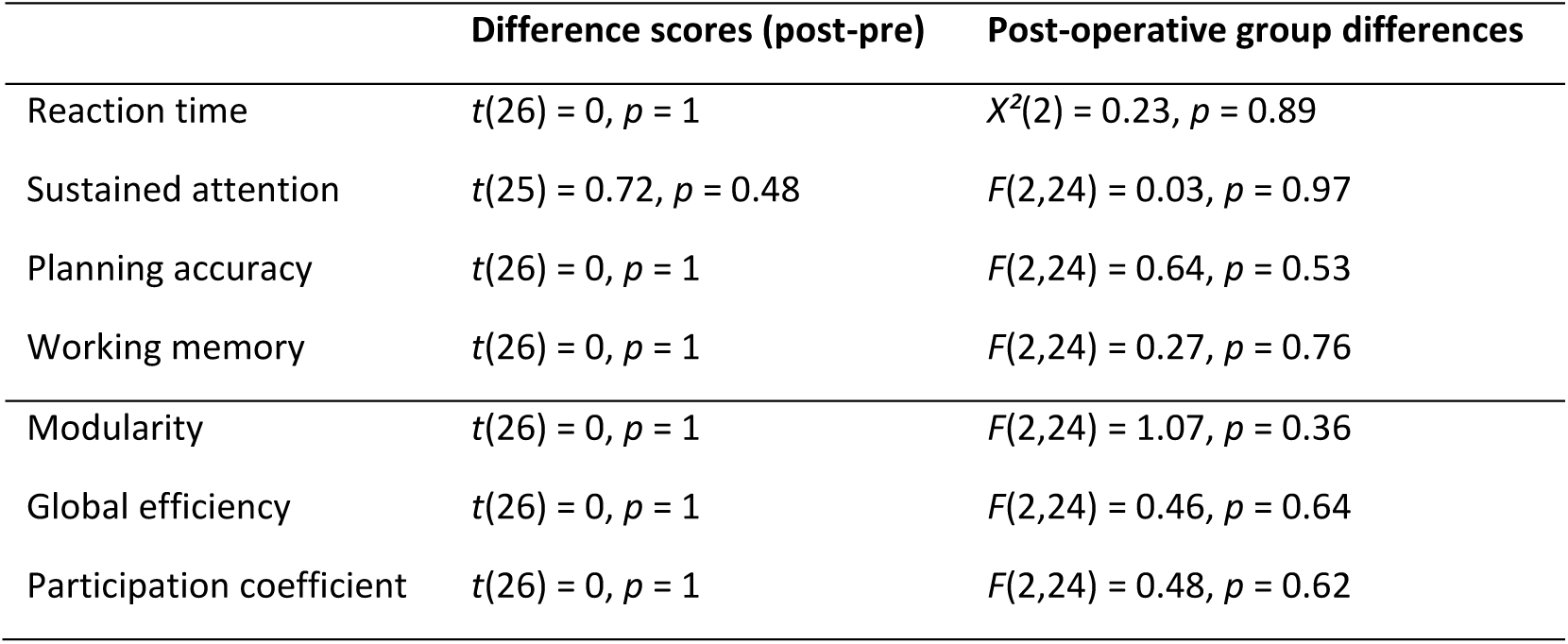
Statistical results of analyses on differences over time and between groups of cognitive performance and structural network topology scores.

|  | Difference scores (post-pre) | Post-operative group differences |
| --- | --- | --- |
| Reaction time | $t(26) = 0, p = 1$ | $\chi^2(2) = 0.23, p = 0.89$ |
| Sustained attention | $t(25) = 0.72, p = 0.48$ | $F(2,24) = 0.03, p = 0.97$ |
| Planning accuracy | $t(26) = 0, p = 1$ | $F(2,24) = 0.64, p = 0.53$ |
| Working memory | $t(26) = 0, p = 1$ | $F(2,24) = 0.27, p = 0.76$ |
| Modularity | $t(26) = 0, p = 1$ | $F(2,24) = 1.07, p = 0.36$ |
| Global efficiency | $t(26) = 0, p = 1$ | $F(2,24) = 0.46, p = 0.64$ |
| Participation coefficient | $t(26) = 0, p = 1$ | $F(2,24) = 0.48, p = 0.62$ |

**Figure 3.**
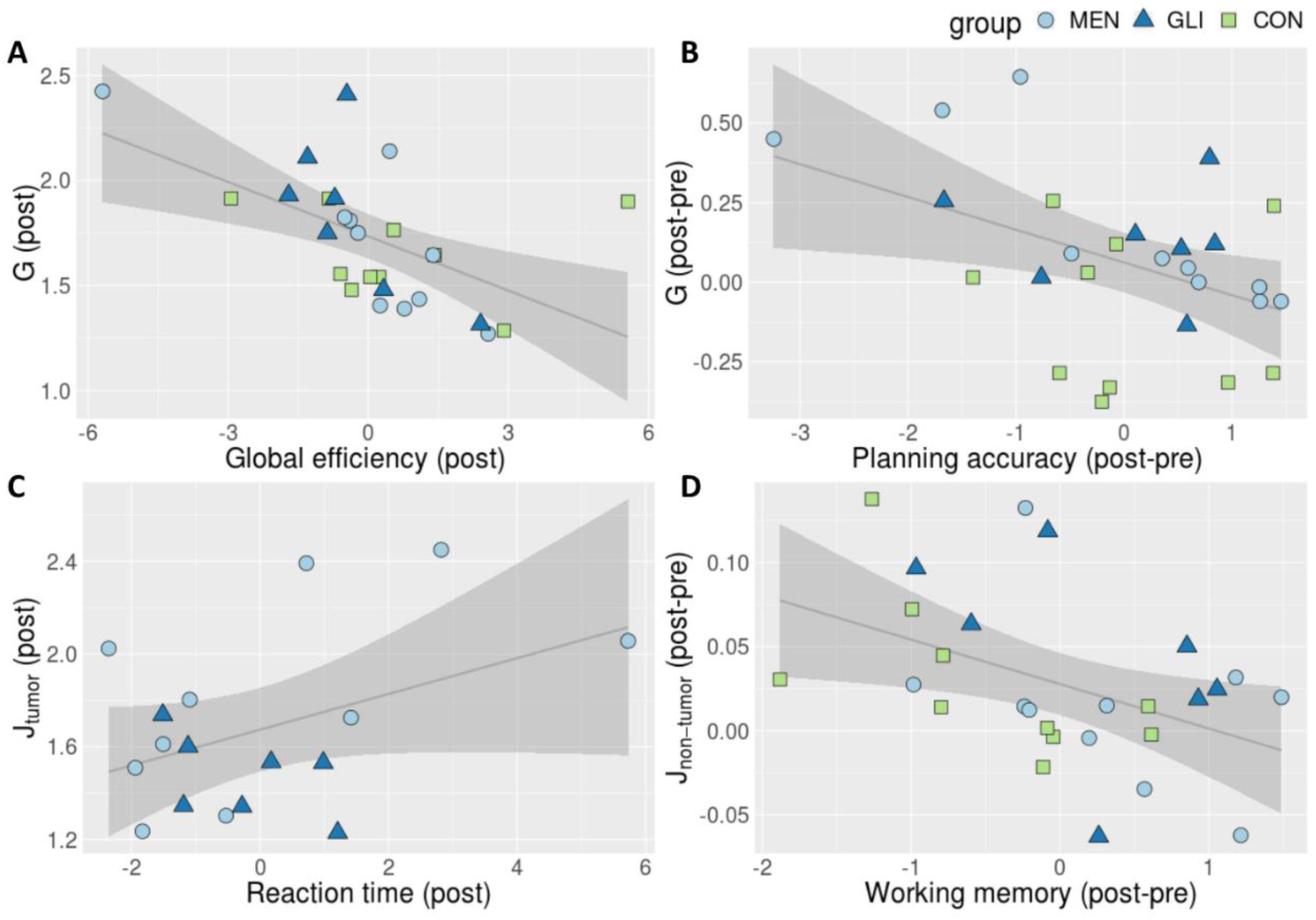
Visual summary of statistically significant linear relationships between individually optimized model parameters, structural network topology and cognitive performance. G = global scaling parameter; Jtumor = median inhibitory synaptic weight across tumor regions in brain tumor patients (corrected for region size); post = calculated on post-operative data; post-pre = difference score between post- and pre-operative data. Gray line represents regression line with 95% confidence interval. Group membership is shape- and color-coded: MEN = meningioma patients; GLI = glioma patients; CON = healthy control participants.

With regard to cognitive performance, results from prior work (Aerts et al., 2018) showed no statistically significant group differences before surgery in any of the cognitive domains assessed. Likewise, we find no significant changes from pre-to post-operative assessment. Consequently, no group differences are apparent in cognitive performance after patients’ tumor resection.

Concerning structural network topology, we find no significant changes over time. Furthermore, no statistically significant group differences are found in these post-operative graph metrics, despite increased levels of participation coefficient in glioma patients before surgery (Aerts et al., 2018).

### 3.3 Robust associations between modeling parameters, structural network topology and cognitive performance

In the next step, we investigate the relations between the individually optimized modeling parameters on the one hand, and structural network topology and cognitive performance on the other hand. Two out of four statistically significant associations that were found pre-operatively could be replicated.

In particular, a negative correlation between global efficiency of the structural network and the global scaling factor is replicated after patients’ tumor resection, as shown in Figure 3A (*t* = −2.77, *p* = 0.0108, *η* ^*2*^ = 0.23; *t* = −2.16, *p* = 0.0420; *η* ^*2*^ = 0.17 after removal of 1 outlier). Likewise, individual differences between pre- and post-surgical optimization of the global scaling parameter are inversely related to the global efficiency of the structural network (*t* = −2.35, *p* = 0.0279, *η* ^*2*^ = 0.19). However, this association is no longer statistically significant after removal of one outlier (*t* = −0.95, *p* = 0.35, *η* ^*2*^ = 0.04).

Next, we find an additional association between pre-to post-surgical differences in the global scaling parameter and planning accuracy (*t* = −2.78, *p* = 0.0113, *η* ^*2*^ = 0.23; *t* = −2.12, *p* = 0.0469, *η* ^*2*^ = 0.15 after removal of 1 outlier; Figure 3B). Although changes in these two variables appear to covary, their association is however not statistically significant before or after surgery (Before surgery: *t* = 0.43, *p* = 0.67; After surgery: *t* = −0.66, *p* = 0.52). In contrast to the results obtained before patients’ surgery, we observed no significant association between participants’ post-operative median inhibitory synaptic weight parameter across healthy regions and the global efficiency of their structural network (*t* = −0.70, *p* = 0.49, *η* ^*2*^ = 0.02).

Furthermore, a significant association is replicated between patients’ median feedback inhibition control parameter across tumor regions and their reaction time during post-operative cognitive assessment (*t* = −2.86, *p* = 0.0144, *η* ^*2*^ = 0.39; Figure 3C). The association between patients’ median feedback inhibition control parameter across tumor regions and their sustained attention is no longer significant (*t* = −1.47, *p* = 0.17, *η* ^*2*^ = 0.10).

Lastly, we observed a statistically significant relation between differences over time in participants’ median inhibitory synaptic weight across healthy brain regions and their working memory capacity (*t* = −2.72, *p* = 0.0127, *η* ^*2*^ = 0.24; Figure 3D).

### 3.4 Virtual neurosurgery proof of concept

In order to evaluate the capacity of the currently applied brain network models to predict patients’ post-surgical brain dynamics, we simulate brain dynamics after virtual neurosurgery and compare the resulting simulated functional connectivity to patients’ empirical post-operative functional connectivity that served as ground truth. As a reference of how well the model can perform for a given patient, we also compute the maximum similarity between each patient’s pre-operative empirical and simulated functional connectome during parameter optimization, without virtual surgery.

Results of these proof of concept analyses are summarized in Figure 4. Compared to the structural connectome that is used as input (SC), simulating functional connectivity by means of brain network modeling (FCsim) usually improves the correspondence with empirically derived functional connectivity. This is also the case during parameter space explorations (PSE: FCsim > PSE: SC). Yet, important individual differences are evident in the extent to which computational modeling can enhance prediction accuracy beyond the structural connectome. As can be seen in Figure 4, substantial improvements in prediction accuracy due to simulating brain activity during parameter space exploration are observed in four out of seven patients, whereas only marginal gains are found in the remaining three patients. Importantly, this aspect – i.e., the degree to which the model can increase prediction accuracy beyond the underlying structure – appears to be of key importance in assessing the potential of this technique for virtual neurosurgery (VS). In particular, prediction of post-surgical brain dynamics only improves in three out of four glioma patients for which computational modeling also yield substantially improved prediction accuracy beyond the structural connectome during parameter space exploration. In the other four patients, correspondence with empirical functional connectivity decreases after simulating virtual neurosurgery compared to using only the virtually lesioned structural connectivity matrix.

**Figure 4.**
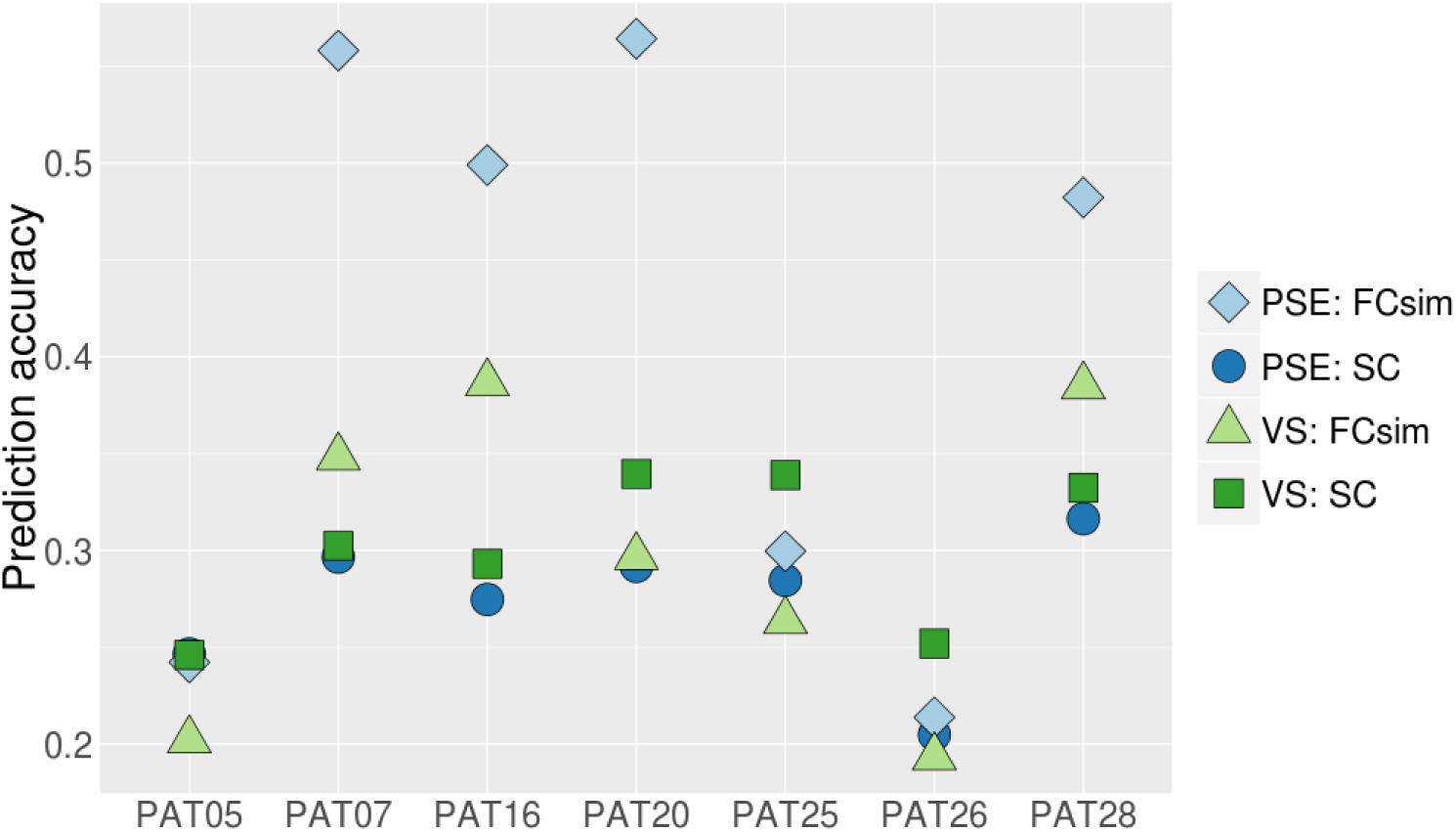
Link-wise Pearson correlation with empirical post-operative FC (as a measure of prediction accuracy) across four different conditions in seven glioma patients that are used in proof of concept analyses for virtual neurosurgery. Specifically, correspondence is evaluated between (1) pre-operative empirical FC and simulated FC after parameter space exploration using pre-operative SC as input (PSE: FCsim); (2) pre-operative empirical FC and pre-operative SC (PSE: SC); (3) post-operative empirical FC and simulated FC after virtual surgery on the SC (VS: FCsim); and (4) post-operative empirical FC and pre-operative SC after virtual surgery (VS: SC).

## 4. Discussion

Results from our study reveal that model parameters describing brain dynamics are relatively stable over time in brain tumor patients who underwent tumor resection, relative to baseline variability levels observed in healthy control subjects. Furthermore, several robust associations between individually optimized model parameters, structural network topology and cognitive performance are identified from pre-to post-operative assessment. Based on these findings, we perform the first proof of concept analyses to evaluate the potential of the currently applied brain network models to predict individual brain dynamics after tumor resection, relying solely on pre-operatively available information. We obtain promising results for a subset of patients, and reveal several limitations and challenges that need to be addressed by future research.

### 4.1 Individual biophysical model parameters and their predictors remain stable over time

In contrast to our expectations, the amount of variability in individually optimized model parameters from pre-to post-operative assessment is comparable between brain tumor patients who underwent neurosurgery and healthy control subjects tested across a similar time interval. This means that there is some variability present over time in brain tumor patients’ optimal model parameters, although there is no systematic trend towards increases or decreases in specific model parameters and the amount of variability does not exceed regular test-retest variability levels in healthy control subjects. Only inhibition control parameters across healthy regions are higher across all participants after surgery compared to pre-operative levels. Of note, we observe similar slight increases in feedback inhibition across tumor regions as well, although these differences do not reach statistical significance given the much larger variability of feedback inhibition values across tumor regions. These elevated levels in feedback inhibition could indicate that more inhibition was required to balance the marginally higher levels of global excitation (i.e., global coupling) after surgery.

In the next step, we evaluate structural network properties and measures of cognitive performance as possible predictors of the individual model parameters. In line with pre-operative results described in prior work (Aerts et al., 2018), initial descriptive analyses show remarkable similarity of cognitive performance and structural network topology across groups as well as over time. Only the increased participation coefficient that was found before surgery in glioma patients is no longer significant after tumor resection, pointing towards a normalization of glioma patients’ structural network topology after surgery. While these findings seem counterintuitive, one other study has also reported comparable structural network topology in brain tumor patients relative to healthy controls (Yu et al., 2016). For cognitive performance, in contrast, the majority of previous studies have reported significant alterations in cognitive functioning as a result of a brain tumor or its subsequent treatment (Klein et al., 2002; Taphoorn & Klein, 2004; Taphoorn et al., 1994; Tucha, Smely, Preier, & Lange, 2000). Possibly, the power of our analyses is not sufficient to detect differences between subjects or changes over time, given the limited sample size. Alternatively, the cognitive tasks (and graph metrics) that we utilize are not sufficiently sensitive to capture these differences.

Despite non-significant group differences or changes over time in cognitive performance and structural network topology, we find several associations between these variables and the individually optimized model parameters. Moreover, we replicate two out of four significant associations that were identified pre-operatively. First, we replicate the inverse relation between global efficiency of the structural network and the global scaling factor after patients’ tumor resection. This implies that higher global coupling values are required in subjects whose structural connectome is less efficiently organized, in order to achieve the same amount of functional connectivity between cortical areas. This appears to be a very robust association, that has also been reported in stroke patients (Falcon et al., 2015). Secondly, we again identify a positive relation between feedback inhibition across tumor regions in brain tumor patients and their reaction time during cognitive assessment. This implies that patients who have higher feedback inhibition take longer to respond on a reaction time assessment. However, similar to results in the pre-operative setting, the association between local inhibitory connection strength and reaction time is largely influenced by a few outlying observations. Hence, caution is advised in interpreting this finding and a larger sample size would be required to clarify this association.

Furthermore, we identify two additional significant associations after patients’ tumor resection. Specifically, differences in the global scaling parameter from pre-to post-surgical assessment are inversely related to changes over time in planning accuracy. Likewise, changes over time in feedback inhibition across healthy regions are negatively associated with differences in working memory capacity from pre-to post-operative assessment. This means that those subjects whose global coupling or feedback inhibition across healthy regions decreased over time, showed improved performance on planning accuracy or working memory tasks, respectively. This finding provides an interesting direction for future research, for example by investigating whether cognitive training can impact model parameters.

### 4.2 Virtual neurosurgery results

Given that brain tumor patients’ model parameters remain relatively stable from pre-to post-operative assessment, we investigate whether post-operative brain dynamics can be predicted using only pre-operatively available information. To this end, we virtually lesion the patient’s pre-operative structural connectome according to the resection mask derived after surgery, based on which we re-simulate brain dynamics using a brain network model with the patients’ pre-surgically optimized global coupling value. Any differences over time in feedback inhibition are assumed not to pose any problems for these proof of concept analyses, as the regional feedback inhibition control parameters are tuned automatically, to clamp the average firing rate at 3 Hz for each excitatory mass model.

Our proof of concept analyses yield promising results for three glioma patients, while the predictive accuracy of the currently applied models is poor in the remaining four patients. Importantly, model performance during pre-operative parameter space exploration appears to be a key indicator for the potential of this technique for virtual neurosurgery. In particular, prediction of post-surgical brain dynamics seems to be feasible only in patients for which the model also substantially improves prediction accuracy beyond the structural connectome during parameter space exploration. Specifically, out of the four patients for which this was the case, prediction accuracy of post-operative brain dynamics is relatively successful in three of them. In one patient simulated post-operative brain dynamics do not show good correspondence with empirical post-operative brain dynamics; possibly the resection mask did not give an appropriate idea of the surgical intervention performed. In the other three patients for which computational modeling only results in marginal gains in prediction accuracy relative to the structural connectome, results show worse prediction accuracy after simulating post-operative brain dynamics compared to using only the virtually lesioned structural connectivity matrix. Nevertheless, for those patients, the virtually lesioned structural connectome serves as a good approximation of their post-operative functional connectivity, suggesting that their brain dynamics are more determined by the underlying structure in the months following tumor resection.

### 4.3 Limitations and future directions

In the interpretation of these study results, some important limitations have to be taken into consideration. First, our sample size is rather small, limiting the statistical power of the analyses. Additionally, substantial inter-subject variability is present in both patient groups, caused by (among other factors) heterogeneity in lesion etiology, size and location. As a result, subtle differences between groups or over time are difficult to detect. By making use of increasingly available open-access clinical datasets, future studies may benefit from using larger sample sizes.

Secondly, although feedback inhibition control parameters are controlled for region size, a substantial association between both remains. This may influence the results, since several tumors overlap with two very large regions (superiorfrontal left and right), whose inhibition values are much higher compared to those of other regions. Currently, model optimization processes are being improved in order to optimize regional feedback inhibition control parameters based on the empirical FC data rather than using the firing rate, which is a direct proxy of region size. Complementary, future research could use parcellation schemes with equally sized regions in order to avoid the confounding effect of region size.

Lastly, important individual differences are observed in the added benefit of simulating brain dynamics on top of the individual structural connectome. These results could reflect true differences in the appropriateness of the model between subjects. Alternatively, however, these differences may result from instabilities in the parameter optimization procedure by maximizing the link-wise Pearson correlation between empirical and simulated FC. Although this method is routinely employed in large-scale modeling studies, other methods that maximize the large-scale organization of both connectivity matrices might be more sensible and yield more robust results. For example, similarity could be sought at the modular level, maximizing the cross-modularity between simulated and empirical functional connectivity (Diez et al., 2015; Stramaglia et al., 2017).

## 5. Conclusion

In summary, our study is the first investigation of potential changes in model parameters describing brain dynamics after brain tumor resection using large-scale brain network modeling. Notwithstanding the methodological caveats described above, we provide preliminary evidence that optimized model parameters are relatively stable from pre-to post-operative assessment. Furthermore, several robust associations between individually optimized model parameters, structural network topology and cognitive performance are identified from pre-to post-operative assessment. Based on these findings, we perform the first proof of concept analyses to evaluate the potential of brain network modeling to predict brain dynamics after tumor resection. We obtain promising results in a subset of patients and reveal important limitations that need to be addressed by future research.

## Acknowledgments

This project has received funding from the Special Research Funds (BOF) of the University of Ghent (01MR0210 and 01J10715), Grant P7/11 from the Interuniversity Attraction Poles Program of the Belgian Federal Government, and the European Union’s Horizon 2020 Framework Programme for Research and Innovation under the Specific Grant Agreement No. 785907 (Human Brain Project SGA2). T.D. acknowledges the National Health and Medical Research Council (NHMRC) of Australia and the Victorian Government’s Operational Infrastructure Support Program for their support. PR acknowledges the following funding sources: H2020 Research and Innovation Action grants 826421 and 650003, 720270 and 785907, and ERC 683049; German Research Foundation CRC 1315 and 936, and RI 2073/6-1; Berlin Institute of Health and Foundation Charité, Johanna Quandt Excellence Initiative. B.J. is a postdoctoral fellow supported by the Research Foundation Flanders (FWO Vlaanderen). Finally, we would like to thank Prof. Dr. Dirk Van Roost, Stephanie Bogaert, Robby De Pauw, Hannes Almgren, Iris Coppieters, Jeroen Kregel, Mireille Augustijn, Helena Verhelst and Kelly Berckmans for their help in acquiring the data.

## Footnotes

1 After the first patient was tested for follow-up, the resting-state fMRI protocol was accidentally changed from a TR of 2100 ms to a TR of 2400 ms resulting in a slightly longer acquisition time of 7:19 minutes. In prior work, we have however demonstrated that such slight alterations in TR have little impact on the construction of FC matrices or the model parameter optimization (Aerts et al., 2018).

