## Supplementary Figure 1 for "Modeling brain dynamics after tumor resection using The Virtual Brain"

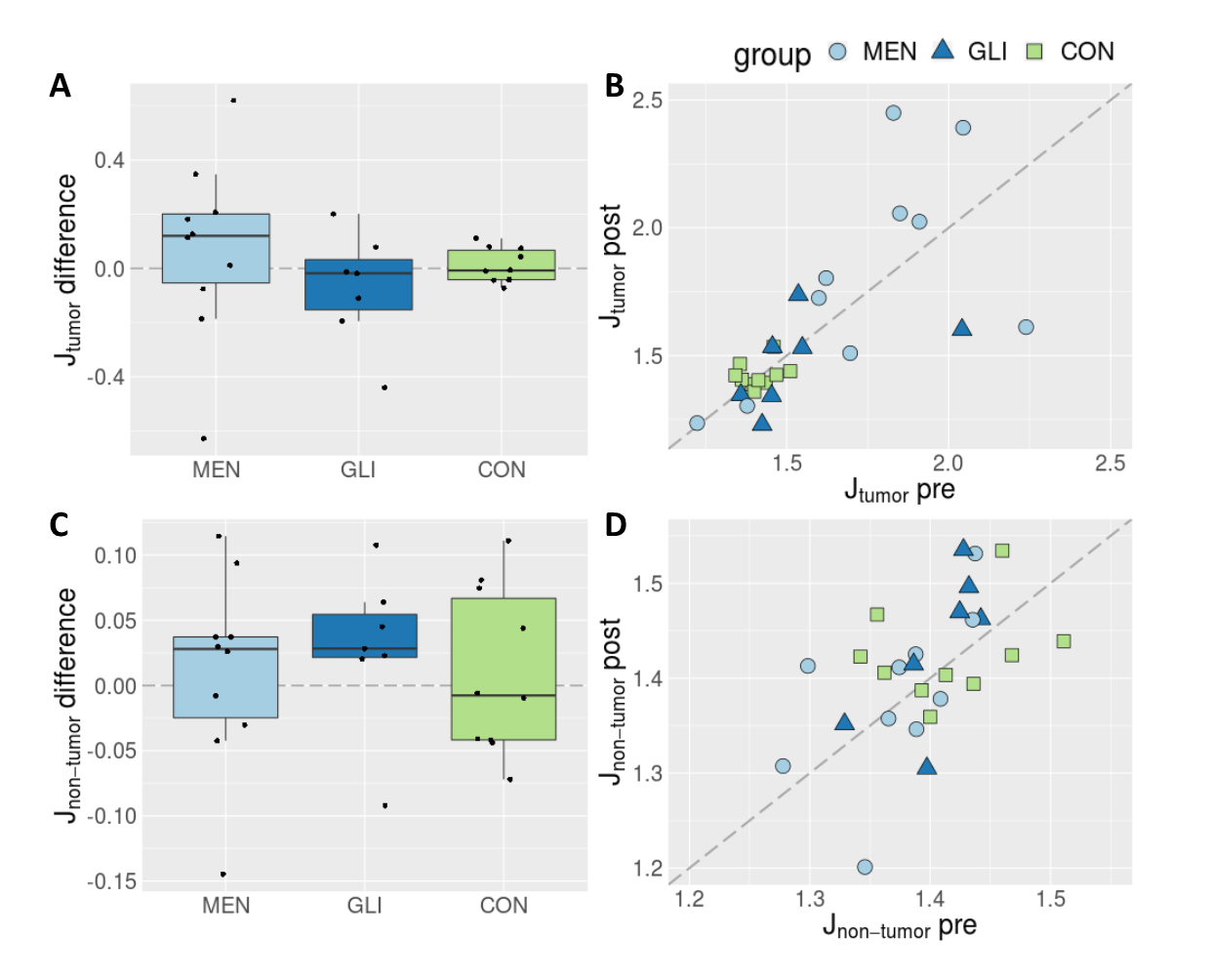

**Supplementary Figure 1.** **Left:** Difference scores between pre- and post-operatively optimized model parameters by group. Differences over time in fitted parameters are shown as deviations from the horizontal line drawn around zero, with positive scores indicating increases in post-operative relative to pre-operative measures, and negative scores corresponding to decreases after surgery compared with pre-operative levels. **Right:** Pre- versus post-operatively optimized individual model parameters, shape- and color-coded by group. Here, differences over time in individuals’ fitted parameters are shown as deviations from the main diagonal, with scores above the diagonal indicating increases in post-operative relative to pre-operative measures, and measures below the main diagonal corresponding to decreases after surgery compared with pre-operative levels. **A & B:** Median local inhibitory connection strengths (not corrected for region size) across tumor regions in meningioma and glioma patients, and across entire brain in healthy controls; **C & D:** Median local inhibitory connection strengths (not corrected for region size) across non-tumor regions in meningioma and glioma patients, and across entire brain in healthy controls. MEN = meningioma patients; GLI = glioma patients, CON = control subjects.
