## Supplementary Figure 2 for "Modeling brain dynamics after tumor resection using The Virtual Brain"

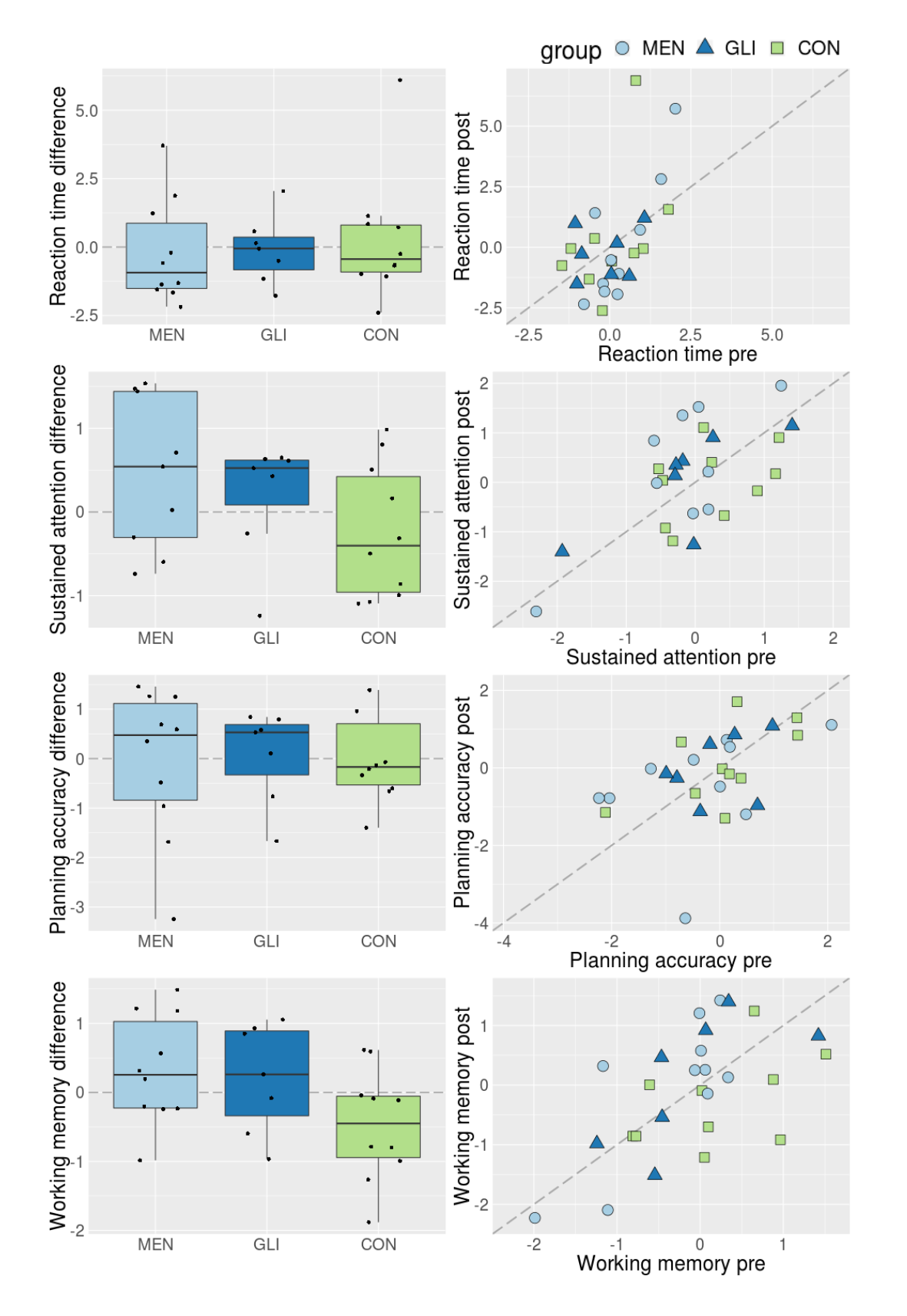

**Supplementary Figure 2.** **Left:** Difference scores between pre- and post-operative cognitive performance measures by group. Differences over time in cognitive performance are shown as deviations from the horizontal line drawn around zero, with positive scores indicating increases in post-operative relative to pre-operative measures, and negative scores corresponding to decreases after surgery compared with pre-operative levels. **Right:** Pre- versus post-operative cognitive performance scores, shape- and color-coded by group. Here, differences over time in individuals’ cognitive performance are shown as deviations from the main diagonal, with scores above the diagonal indicating increases in post-operative relative to pre-operative measures, and observations below the main diagonal corresponding to decreases after surgery compared with pre-operative levels. All metrics are corrected for important confounding variables (described in section 2.6 “Accounting for covariates of no immediate interest”) and transformed to z-scores using the pre-operative mean and standard deviation of the respective metric in the group of control subjects. MEN = meningioma patients; GLI = glioma patients, CON = control subjects.
