## Supplementary Figure 3 for "Modeling brain dynamics after tumor resection using The Virtual Brain"

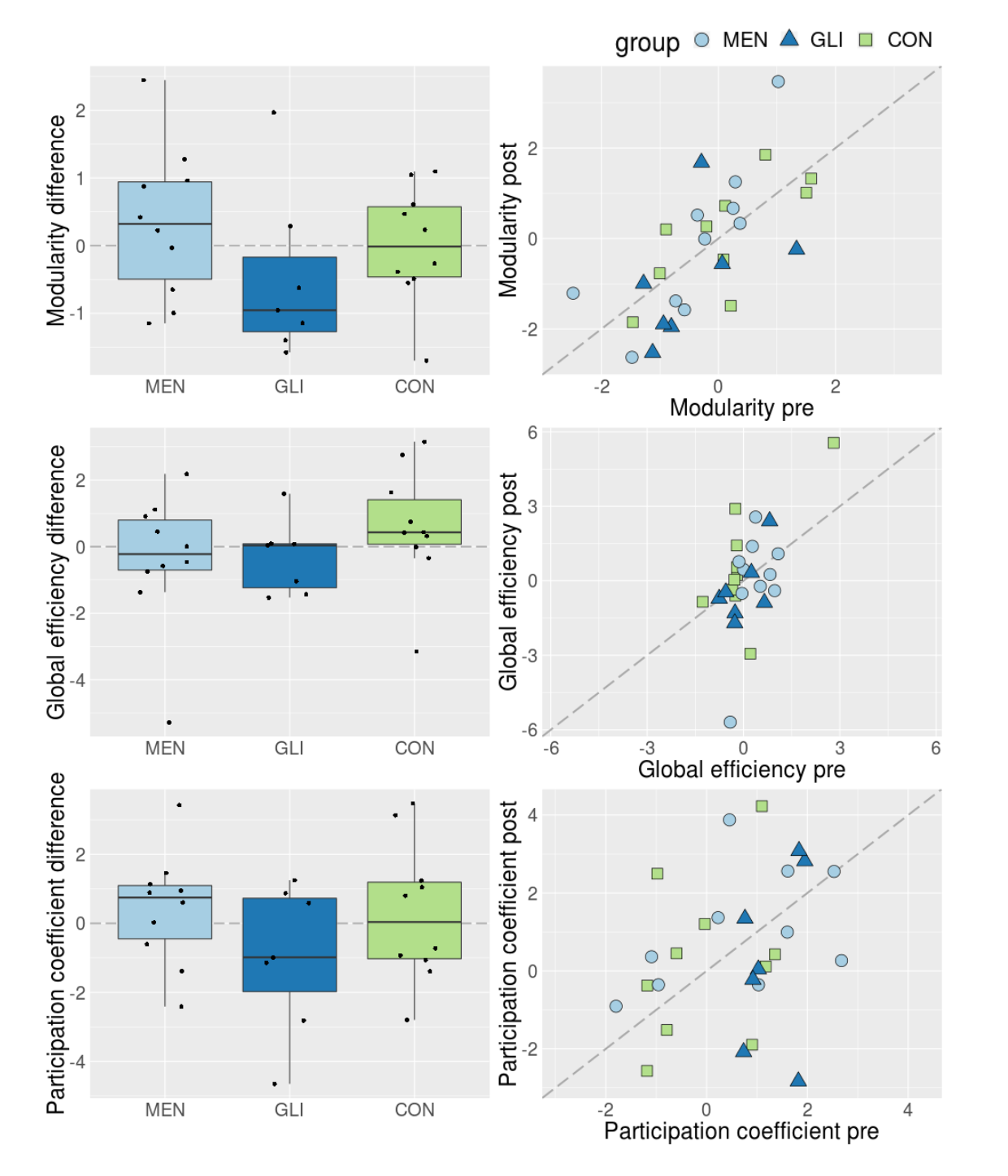

**Supplementary Figure 3.** **Left:** Difference scores between pre- and post-operative structural network topology measures by group. Differences over time in graph measures are shown as deviations from the horizontal line drawn around zero, with positive scores indicating increases in post-operative relative to pre-operative measures, and negative scores corresponding to decreases after surgery compared with pre-operative levels. **Right:** Pre- versus post-operative structural network topology scores, shape- and color-coded by group. Here, differences over time in individuals’ cognitive performance are shown as deviations from the main diagonal, with scores above the diagonal indicating increases in post-operative relative to pre-operative measures, and observations below the main diagonal corresponding to decreases after surgery compared with pre-operative levels. All metrics are corrected for important confounding variables (described in section 2.6 “Accounting for covariates of no immediate interest”) and transformed to z-scores using the pre-operative mean and standard deviation of the respective metric in the group of control subjects. MEN = meningioma patients; GLI = glioma patients, CON = control subjects.
